# Effects of four antibiotics on the diversity of the intestinal microbiota

**DOI:** 10.1101/2021.07.19.453004

**Authors:** Ce Huang, Shengyu Feng, Fengjiao Huo, Hailiang Liu

## Abstract

Oral antibiotics remain the therapy of choice for severe bacterial infections; however, antibiotic use disrupts the intestinal microbiota, which increases the risk of colonization with intestinal pathogens. Currently, our understanding of antibiotic-mediated disturbances of the microbiota remains at the level of bacterial families or specific species, and little is known about the effect of antibiotics on potentially beneficial and potentially pathogenic bacteria under conditions of gut microbiota dysbiosis. Additionally, it is controversial whether the effects of antibiotics on the gut microbiota are temporary or permanent. In this study, we used 16S rRNA gene sequencing to evaluate the short-term and long-term effects of ampicillin, vancomycin, metronidazole, and neomycin on the murine intestinal microbiota by analyzing changes in the relative numbers of potentially beneficial and potentially pathogenic bacteria. We found that the changes in the intestinal microbiota reflected the antibiotics’ mechanisms of action and that dysbiosis of the intestinal microbiota led to competition between the different bacterial communities. Thus, destruction of bacteria with beneficial potential increased the abundance of bacteria with pathogenic potential. In addition, we found that these oral antibiotics had long-term negative effects on the intestinal microbiota and promoted the development of antibiotic-resistant bacterial strains. These results indicate that ampicillin, vancomycin, metronidazole, and neomycin have long-term negative effects and can cause irreversible changes in the diversity of the intestinal microbiota and the relative proportions of bacteria with beneficial potential and bacteria with pathogenic potential, thereby increasing the risk of host disease.

## INTRODUCTION

The host intestinal tract provides the intestinal microbiota with an anaerobic or hypoxic nutrient-rich environment, and as a result hundreds of millions of different types of microorganisms parasitize the human intestine, forming a complex ecological environment (Sommer et al., 2017). Microorganisms within the intestinal microbiota can be divided into three categories based on their interaction with the host: mutualists (benefiting themselves and the host), commensals (benefiting themselves but not the host), and pathogens (benefiting themselves by harming the host) (Reid et al., 2011). Here we will refer to mutualists as “potentially beneficial bacteria”. Potentially beneficial bacteria play a very important role in maintaining the stability of the intestinal microbiota and also provide the host with biochemical metabolic pathways and enzymes that it does not already have, help the host to break down nutrients, synthesize nutrients needed by the human body, maintain the stability of the nervous system, activate the human immune system, and maintain intestinal homeostasis (Kim et al., 2013; Sanders et al., 2019). For example, *Turicibacter, Coprobacillus*, and *Lactobacillus* produce lactic acid through glycolysis, which can provide intestinal epithelial cells with nutrition and regulate immune function (Getachew et al., 2018; Liévin-Le Moal and Servin, 2014; Muthuramalingam et al., 2020), and *Parabacteroides, Faecalibacterium, Odoribacter* and *Coprococcus* can reduce intestinal inflammation by secreting acetic acid and butyrate, thereby decreasing the incidence of inflammatory colitis, Crohn’s disease and other diseases (Ferreira-Halder et al., 2017; González-Sarrías et al., 2018; Han et al., 2020; Wang et al., 2019). *Roseburia* and *Akkermansia* can ferment a variety of carbohydrates and have been used to treat diseases such as obesity and diabetes (Depommier et al., 2019; Seo et al., 2020). Pathogenic bacteria account for a small proportion of the microorganisms in the intestine. Many of these species, when they are present in small quantities, are an important part of the healthy intestinal microbiota. However, when intestinal microbiota dysbiosis occurs, these pathogens will harm the host. Among them, *Enterococcus, Enterobacter*, and *Clostridium perfringens* are the main pathogenic bacteria. They break down some components of food into amines through their own unique biochemical metabolic pathways. However, they also produce indole, phenols, and unique microbial toxins, which cause certain intestinal diseases (Bäumler and Sperandio, 2016; Perez-Lopez et al., 2016). For example, *Clostridium difficile* growth and reproduction can cause diarrhea and pseudomeningitis (Smits et al., 2016), and an excessive number of enterococci can cause abdominal and pelvic infections (Arias and Murray, 2012). When the number of potentially beneficial bacteria in the human intestine decreases and the number of pathogenic bacteria increases, intestinal homeostasis is disrupted, which can cause host metabolic disorders and affect the host’s immune system.

Antibiotics are frequently used to treat bacterial infections. Epidemiological studies have shown that 71% of Intensive Care Unit (ICU) patients are on antibiotics. Long-term, excessive use of antibiotics can cause serious adverse consequences. Although antibiotics inhibit the growth of and kill bacteria, they can also induce drug resistance. In addition, antibiotic use can temporarily or permanently alter the composition of the intestinal microbiota, promote colonization with intestinal pathogens, and trigger the development of some intestinal diseases (Freifeld et al., 2011; Gilbert et al., 2012). In recent years, disruption of the intestinal microbiota by antibiotics has received increasing attention. Some studies have demonstrated a direct link between alterations in the intestinal microbiota and antibiotic use. For example, short-term treatment of mice with metronidazole, vancomycin and clindamycin can reduce their susceptibility to infection with *C. difficile*, vancomycin-resistant *Enterococci, Klebsiella pneumonia*, and *Escherichia coli* (Buffie et al., 2012; Lewis et al., 2015). In this study we investigated the effects of four commonly used antibiotics: Vancomycin, which is typically used to treat infections with Gram-positive bacteria, Ampicillin and Neomycin, which are typically used for Gram-negative bacterial infections, and Metronidazole, which is typically used to treat anaerobic bacterial infections, to account for most of the bacterial components of the intestinal microbiota. To date, studies of the relationship between these antibiotics and the intestinal microbiota have mostly focused on the impact of antibiotics on colonization with potentially pathogenic bacteria such as *C. difficile* and *Enterobacter* (Lewis et al., 2015). In addition, these studies of microbiota disruption associated with antibiotic use have remained at the level of bacterial families or specific species (Dethlefsen et al., 2008; Rodrigues et al., 2017; Sun et al., 2019). Combined treatment with ampicillin, vancomycin, neomycin, and metronidazol provides bactericidal activity against the full spectrum of bacteria and, notably, dual activity against both Gram-positive (ampicillin and vancomycin) and Gram-negative (ampicillin and neomycin) aerobic and facultative strains (Table S1). Few studies have used high-throughput 16s rRNA gene sequencing technology to conduct in-depth analysis of changes in intestinal microbiota diversity in response to these antibiotics treatment, and especially to quantitatively assess the relative proportions of potentially beneficial bacteria and potentially pathogenic bacteria.

On the other hand, a number of recent studies have reported the use of a combination of four antibiotics added to drinking water to eliminate the gut microbiota, reducing the fecal DNA content to 3% of the control (Le Roy et al., 2018; Miao et al., 2020; Yang et al., 2017; Zhang et al., 2020). However, most of these studies only quantified the total DNA content of the feces, which demonstrated that the gut microbiota was reduced, but did not provide any detailed information regarding the effect of this treatment regimen on intestinal microbiota dysbiosis. There is also a lack of research on whether this combination of antibiotics can minimize the diversity of intestinal microbiota compared with single antibiotic use.

In this study, we used 16S rRNA gene sequencing technology to analyze the short-term and long-term effects of ampicillin, vancomycin, metronidazole, and neomycin on the murine intestinal microbiota by assessing changes in the relative numbers of bacteria with beneficial potential and bacteria with pathogenic potential. In addition, we compared a mixture of four antibiotics with a single antibiotic to analyze the effect of this mixture on the diversity of the intestinal flora. Our findings provide clinically relevant information for the treatment of bacterial infections and verify the effectiveness of current methods that are commonly used to eliminate the intestinal microbiota.

## MATERIALS AND METHODS

### Antibiotics

Ampicillin (A5354), Vancomycin (V0045000), Metronidazole (M3761), and Neo mycin (33492) were purchased from Sigma-Aldrich (St. Louis., MO, USA).

### Animals

Eight-week-old C57BL/6 mice were obtained from Shanghai SLAC Laboratory Animal Co., Ltd. Mice (N = 5 per group) were maintained in an SPF room in the Experimental Animal Center at Tongji University, in cages with free access to water and food and a 12-hour/12-hour light/dark cycle. After the mice had been acclimatized for 10 weeks, one mouse per cage. We have a total of 24 cage mice, numbered sequentially, and randomly generated Numbers up to 24 using the Microsoft Excel Randbetween function. Three Numbers are grouped into groups. And antibiotics were added to the drinking water based on the weight of the mice. There were six short-term experimental groups (male mice): the control group; the AMP group (100 mg/kg Ampicillin); the Van group (50 mg/kg Vancomycin); the Met group (100 mg/kg Metronidazole); the Neo group (100 mg/kg Neomycin); and the S-Mix group (100 mg/kg Ampicillin; 50 mg/kg Vancomycin; 100 mg/kg Metronidazole; and 100 mg/kg Neomycin). In addition, there were two long-term experimental groups (male mice): the control group and the L-Mix group (100 mg/kg Ampicillin;50 mg/kg Vancomycin;100 mg/kg Metronidazole;100 mg/kg Neomycin). The AMP, Van, Met, Neo, and S-Mix groups received antibiotics for 14 consecutive days, followed by a 30-day washout period, followed by antibiotic administration for another 14 days. The L-Mix group received antibiotics for 60 days. The ethics committee of Tongji University approved all of the protocols used in this study. After an overnight fast at the end of the feeding period, mice were deeply anaesthetized using tribromoethanol (Sigma, St. Louis, MO). Blood was taken by removalling eyeball and centrifuged at 1000g/min for 10min at 4℃ to obtain plasma. After the experiment, we treated the mice with physical euthanasia, the mice were euthanized by cervical dislocation. There were no adverse events.

### Genomic DNA extraction

Total microbial DNA was extracted using a MagPure stool DNA KF kit B (Magen, China) according to the manufacturer’s instructions. DNA was quantified with a Qubit Fluorometer using a Qubit dsDNA BR Assay kit (Invitrogen, USA), and the quality was checked by running an aliquot on a 1% agarose gel.

### Library Construction

The V4 variable region of the bacterial 16s rRNA gene was amplified with degenerate PCR primers: 515F (5′-GTGCCAGCMGCCGCGGTAA-3′) and 806R (5′-GGACTACHVGGGTWTCTAAT-3′). Both primers were tagged with Illumina adapter, pad, and linker sequences. PCR enrichment was performed in a 50-uL reaction containing 30 ng template, both tagged PCR primers, and PCR master mix. The PCR cycling conditions were as follows: 95°C for 3 minutes; 30 cycles of 95°C for 45 seconds, 56°C for 45 seconds, and 72°C for 45 seconds; and a final extension for 10 minutes at 72°C. The PCR products were purified using Agencourt AMPure XP beads and eluted in elution buffer. Libraries were evaluated using an Agilent Technologies 2100 bioanalyzer. The validated libraries were then sequenced on an Illumina HiSeq 2500 platform (BGL, Shenzhen, China) following standard Illumina protocols and generating 2 × 250 bp paired-end reads.

### Bioinformatics analysis

First, low quality was removed from the original sequencing data by the window method. Joint pollution reads were removed, n-containing reads were removed and low-complexity reads were processed. Samples were distinguished based on barcode and primer. After, use FLASH software (fast Length adjustment of short reads, v1.2.11)(Magoč and Salzberg, 2011). Using overlapping relationship will be pairs of double end sequencing reads assembled into a sequence with high area tags. Effective tags were produced by the UCHIME algorithm and clustered into operational taxonomic units (OTUs) with USEARCH (V7.0.1090) software (Edgar, 2013). According to Mothur method and GreenGene database, taxonomic information was annotated with representative sequences from OTU (Schloss et al., 2009). Through R software (v3.4.1, R Core Team (2018)) sample analyzing the complexity of species, species differences between group analysis, correlation analysis and model forecast. Through Kruskal-Wallis test and Wilcoxon test analyzing differential species.

## RESULTS

### Treatment with four antibiotics reduced the richness and diversity of the intestinal microbiota in mice

As shown in Figure 1A, eight groups of three mice were investigated. Antibiotic concentrations were based on the weight of the mice, and the antibiotics were continuously administered for 14 days. We collected fecal samples from each group before the start of treatment and on the 5th and 15th days after treatment began, and performed 16S rRNA-seq analysis on DNA isolated from the collected fecal material. First, we carried out a unweighted pair-group method with arithmetic means (UPGMA) cluster analysis to divide the samples into groups based on microbiota composition. Samples with high similarity in terms of microbiota composition were clustered into the same branch of the evolutionary tree. As shown in Figure 1B, the baseline samples for all of the groups clustered together. After antibiotic treatment was started, the microbiota composition of each group began to change and gradually diverge from the respective baseline samples. Subsequently, we analyzed the impact of the four antibiotic treatments on the diversity of the intestinal microbiota as measured by the number of identified OTUs, as shown in Figure 1C. The curves gradually fell, indicating that the richness of the microbial diversity was greatly reduced. Alpha diversity analyses assess species diversity within a single sample, and include the observed species index, the Chao index, the Ace index, the Shannon index, the Simpson index, and the Goodcoverage index. To perform an alpha diversity analysis, we selected the Shannon index. There was no significant difference in the mean Shannon diversity index before and after antibiotic treatment (Figure 1D; P > 0.05), but the microbial richness of the Van, Met, Neo, and S-Mix groups decreased compared with the control group. Beta diversity analysis (Figure 1E) showed that the microbial richness in the Amp (P<0.01), Van (P<0.05), and S-Mix (P<0.05) groups decreased compared with the control group. The antibiotics used to treat these three groups are active against Gram-negative bacteria, Gram-positive, and both Gram-negative and Gram-positive bacteria, respectively. Neomycin and metronidazole target aerobic and anaerobic bacteria, respectively, but they had no significant effect on the diversity of intestinal bacteria. In terms of the diversity of the gut microbiota, the combination of four antibiotics was inferior to ampicillin or vancomycin alone. This indicates that the combination of multiple antibiotics did not have the greatest impact on intestinal flora, and that the combination of these four antibiotics is not necessarily the optimal choice for eliminating the intestinal microbiota in experimental mouse models.

**Figure 1.**
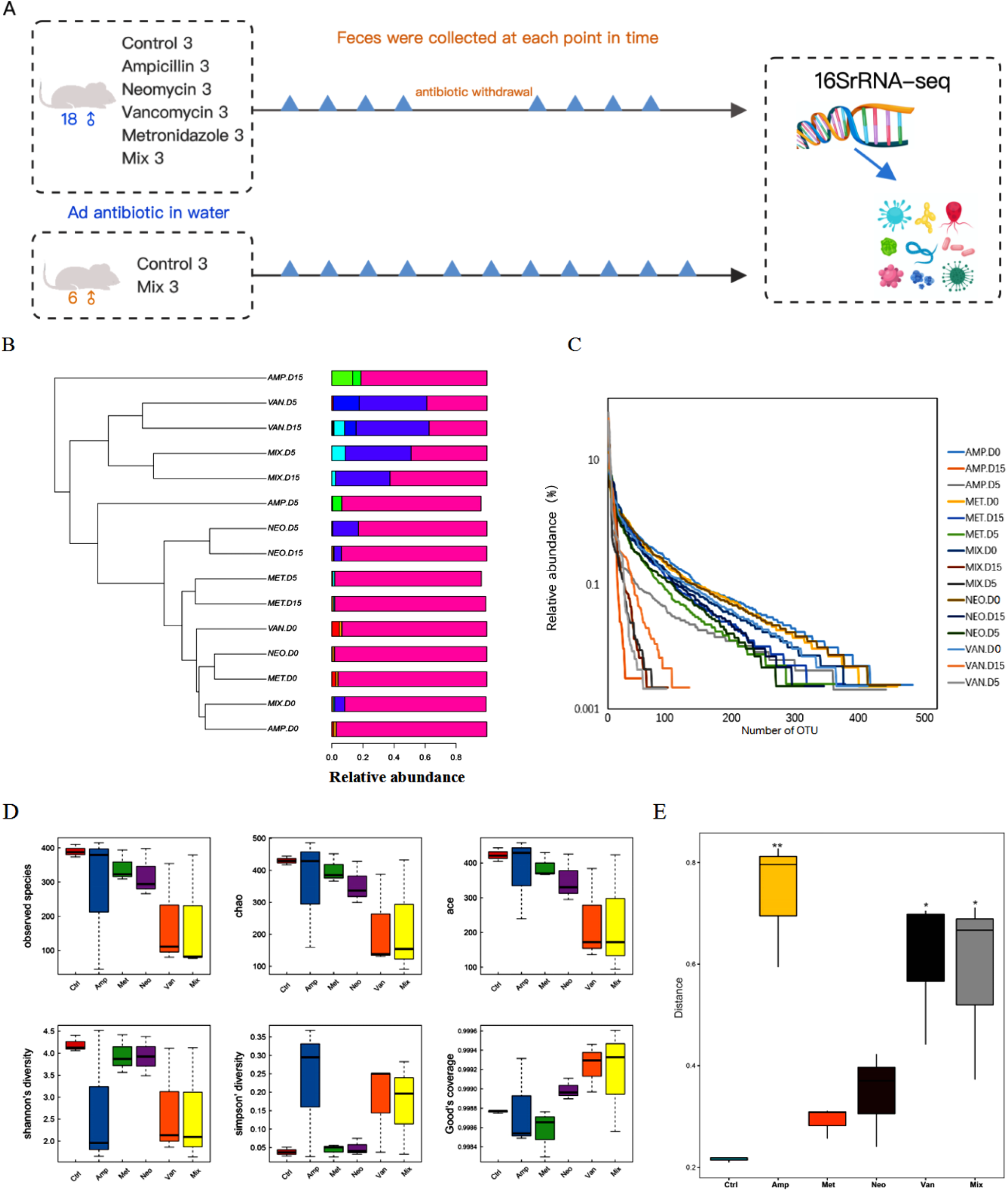
Continuous treatment with four antibiotics alters the diversity an d abundance of the intestinal microbiota in mice. (A) This experiment included eight groups of three 12-month-old mice. The groups were treated with 10 0 mg/kg Ampicillin, 50 mg/kg Vancomycin, 100 mg/kg Metronidazole, or 100 mg/kg Neomycin. Fecal samples were collected at eight time points for the short-term experimental groups and at 11 time points for the long-term experimental groups. (B) Met: Metronidazole, Amp: Ampicillin, Van: Vancomycin, Neo: Neomycin, Mix: Mixture. Microbial UPGMA clustering for fecal samples taken at different time points. Similar samples cluster together on the same branch of the tree. D0 indicates baseline, and D indicates different time points. (C) OTU rank curve. The abscissa is sorted by OTU abundance in each sample (high to low), and OTU abundance is on the ordinate axis. The richness of spec ies within the sample is reflected by the horizontal spread of the curve: the wider the curve, the richer the species composition of the sample. (D) Alpha diversity analysis of the microbiota after 14 days of antibiotic treatment. Alpha diversity analysis assesses species diversity within a single sample, and includes the observed species index, the Chao index, the Ace index, the Shannon index, the Simpson index, the Goodcoverage index, and more. (E) Beta diversity.

### Effects of four antibiotics on potentially beneficial bacteria and potentially pathogenic bacteria in the murine intestinal microbiota

Considering the substantial impact of antibiotics on the richness and diversity o f the intestinal microbiota, we next decided to study its impact on specific tax a. We compared the treatment group and the control group with matched pair t -test on phylum and genus level (Table S2, Table S3). And as Figure 2A and Figure 2B show that *Proteobacteria* increased after Ampicillin treatment at the *phylum* level, *TM7* and *Firmicutes* decreased and the bacteria with beneficial p otential *Ruminocuccus, Coprococcus* of the *Firmicutes* decreased at the genus level, while bacteria with pathogenic potential *Proteus* in *Proteobacteria* and *Bacteroides* in *Bacteroidetes* increased. *Verrucomicrobia* and *Proteobacteria* increased after Vancomycin treatment at the phylum level, *Actinobacteria* (p<0.05), an d *TM7* (p<0.05) decreased, *Verrucomicrobia* (p<0.05) increased and at the genus level, the bacteria with beneficial potential *Oscilillospira* (p<0.05), *Allobaculum* and *Coprococcus* of the *Firmicutes* decreased, while the bacteria with pathogenic potential *Proteus* in *Proteobacteria* and *Bacteroides* in *Bacteroidetes* increased. After treatment with Metronidazole, *Actinobacteria, Bacteroidetes* and *Cyanobacteria* (p<0.05) at the phylum level increased, *Proteobacteria, Firmicutes decreased* at the genus level, the bacteria with beneficial potential *Coprococcus* o f the *Firmicutes* decreased, while the bacteria with pathogenic potential *Sutterella* in *Proteobacteria*, the *Prevotella* and *Bacteroides* increased. After treatment with Neomycin, *Verrucomicrobia* increased while TM7 and *Proteobacteria* (p<0. 05) decreased at the phylum level, it is worth noting that the bacteria with beneficial potential *Allobaculum* and *Coprococcus* (p<0.05) of the *Firmicutes* have increased. After treatment with Mixture, *Actinobacteria* (p<0.05) and *Verrucom icrobi* increased at the phylum level, *Cyanobacteria, TM7* (p<0.05), *Firmicutes* as well as *Proteobacteria* decreased. Also, the bacteria with beneficial potential *Ruminocuccus, Allobaculum* and *Coprococcus* of the *Firmicutes* decreased at t he genus level, while the bacteria with pathogenic potential *Sutterella* and *Bacteroides* of the *Proteobacteria* decreased.

**Figure 2.**
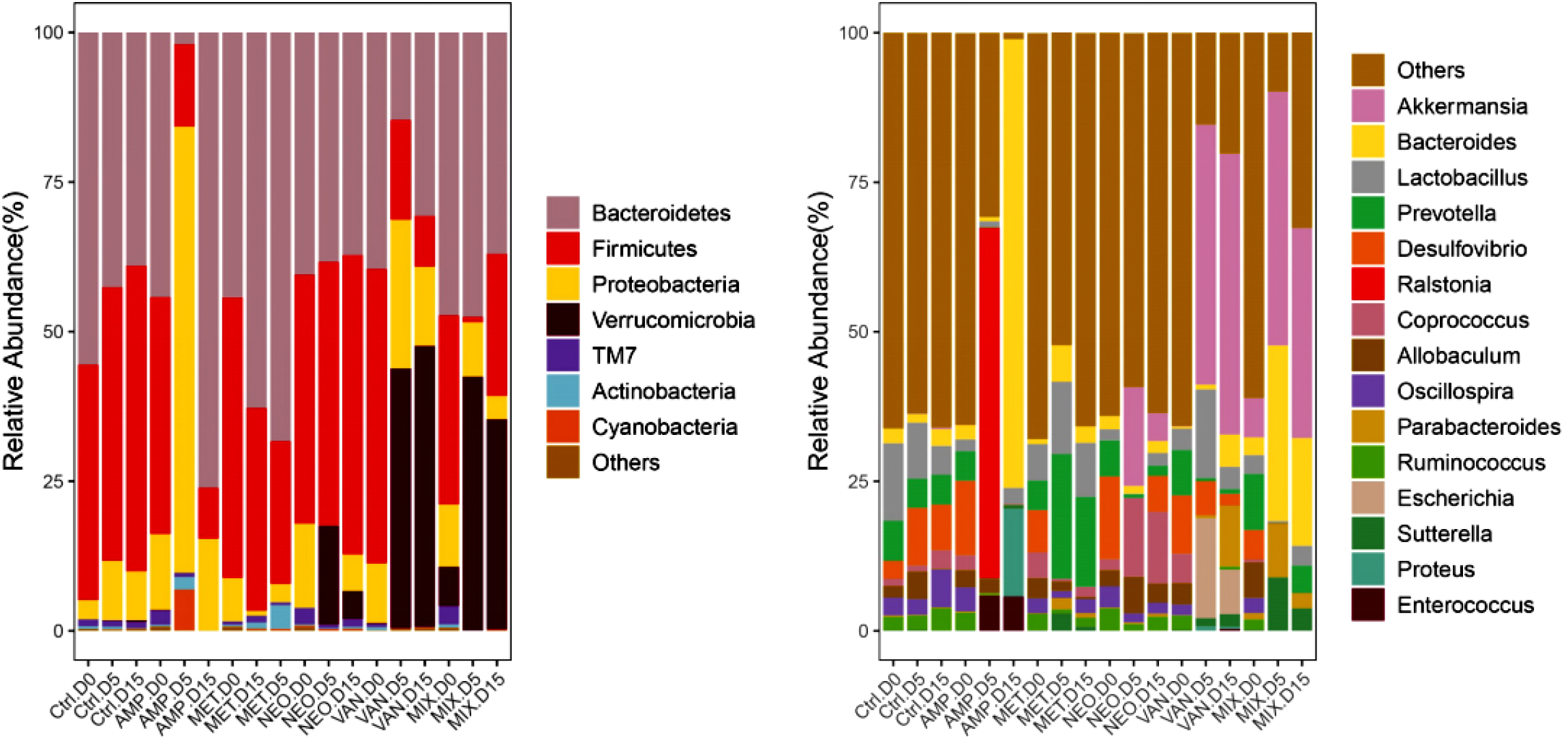
Treatment with the four antibiotics altered the abundance of most taxa. Histograms of species abundance. The sample names are on the abscissa, and the relative abundance of the annotated species is shown on the ordinate axis. Species with less than 0.5% abundance in all samples were merged into the ‘Others’ category. (A) shows abundance at the phylum level, and (B) shows abundance at the genus level.

In general, the balance between bacteria with beneficial potential and bacteria with pathogenic potential in the intestinal microbiota was disrupted by the use of antibiotics. However, different antibiotics have different effects on the two types of intestinal microbiota. *Coprococcus* and *Lactobacillus* are two types of Gram-positive anaerobic bacteria with beneficial potential. Amp, Met, and Van all reduced the abundance of these two bacteria. On the other hand, *Bacteroides*, a Gram-negative anaerobic bacterium with pathogenic potential, should theoretically have been killed by Amp and Met; however, treatment with Amp led to an increase in Bacteroides abundance, and Met had no effect on its abundance. In addition, we found that the abundance of all bacterial communities was not reduced in the Mix group, and treatment with Amp alone reduced the abundance of most of the bacterial communities. These findings suggest that the changes that we observed in the intestinal microbiota reflect not only the mechanisms of action of the antibiotics (Table S3), but also competition between different bacterial communities induced by intestinal microbiota dysbiosis. For example, destruction of bacteria with beneficial potential enabled bacteria with pathogenic potential that were not affected by the relevant antibiotic to increase in abundance. Our results show that there was a consistent reduction in the abundance of bacteria with beneficial potential and a consistent increase in the abundance of the bacteria with pathogenic potential, which could reduce the host’s ability to resist inflammation and increase the risk of infection.

### Recovery of the murine intestinal microbiota after antibiotic withdrawal

Next, we investigated the ability of the murine intestinal microbiota to recover 1 month after antibiotics were withdrawn. We focused on those bacteria whose relative abundance changed significantly during antibiotic treatment. As shown in Figure 3A, 1 month after stopping antibiotic therapy, the composition of the intestinal microbiota in mice from the Amp, Van, Neo, and S-Mix groups did not return to baseline at the OTU level; only the Met group returned to baseline. At the phylum level, as shown in Figure 3B, only the abundance of *Actinobacteria* (0.0028%, 6.23E-05%, 0.0005%) (D0, D15, D45) in the Ampicillin group and *Proteobacteria* (0.1038%, 0.03923%, 0.0881%), *Bacteroidetes* (0.4728%, 0,3698%, 0.4986%), *Firmicutes* (0.3159%, 0.2374%, 0.3306%), and *Verrucomicrobia* (0.0657%, 0.3511%, 0.0810%) in the Mixture group returned to baseline levels, while the previously detected *Actinobacteria* and *Cyanobacteria* phyla disappeared. The abundance of only a few genera fully recovered. As shown in Figure 3C, the abundance of *Ruminocuccu*s (0.0292%, 3.12E-05%, 0.0160%), which had previously decreased, was restored to baseline levels in the Ampicillin group; in the Vancomycin group, the abundance of the bacteria with beneficial potential *Oscillospira* (0.0184%, 4.50E-05%, 0.0363%) and *Coprococcus* (0.0493%, 6.75E-05%, 0%) recovered. *Parabacteroides* and *Escherichia*, which had not previously reached the detection limit in the Neo group, were detected 1 month after neomycin treatment was stopped. The abundance of *Bacteroides* (0.0297%, 0.1801%, 0.0218%), which had decreased, and *Allobaculum* (0.0591%, 2.24E-05%, 0.0792%) and *Coprococcus* (0.0050%, 2.24E-05%, 0.0046%), which had increased, returned to baseline levels in the Mixture group. In addition, the abundance of the *Proteus* and *Enterococcus* genera, which had been detected prior to antibiotic treatment, did not reach the detection limit, and *Ralstonia*, which did not appear prior to treatment, was detected 1 month after withdrawal of the antibiotics.

**Figure 3.**
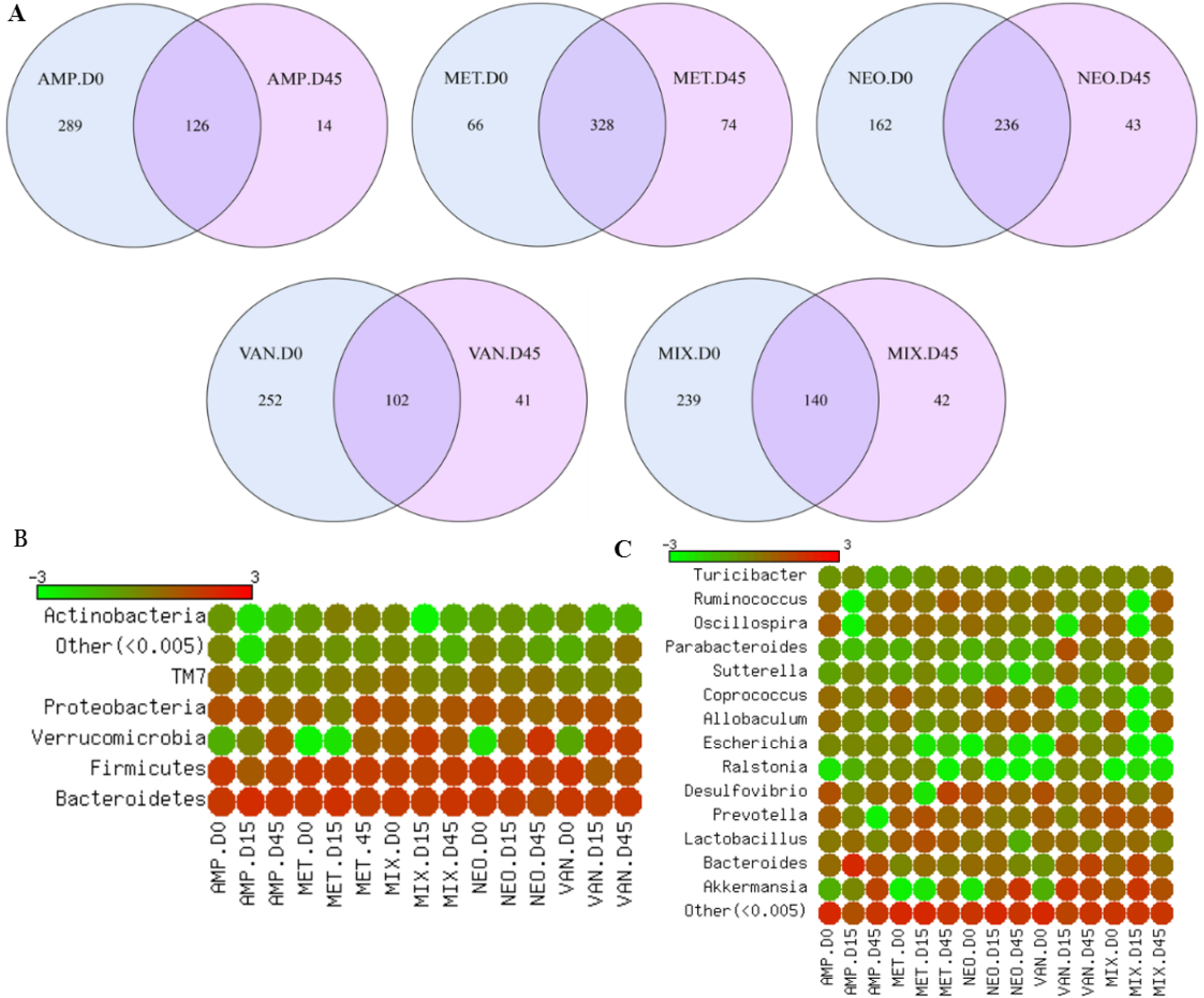
Incomplete and antibiotic-specific recovery of the murine intestinal microbiota after antibiotic withdrawal. (A) The differently colored circles in the Venn diagrams represent different samples or groups. The numbers shown in the overlapping areas indicate the number of OTUs shared between the two samples or groups. The heat maps show the average fold changes and significant increases (red) or decreases (green) (P < 0.05, FDR < 0.2, two-tailed Wilcoxon test) in the abundance of specific phyla (B) and genera (C) compared with baseline. Species whose abundance was less than 0.5% in all samples were merged into the ‘Others’ category.

After antibiotic treatment, the abundance of some bacteria in the intestinal microbiota returned to their baseline levels. However, the abundance of most of the microbiota components was altered, in some cases permanently, and the microbiota composition after antibiotic withdrawal was different from that observed before treatment. Our findings show that, while the abundance of some intestinal microbiota components was restored to pretreatment levels, that of most of the intestinal microbiota components did not recover, and instead remained elevated. The microbiota composition of groups that were treated with a single antibiotic exhibited less recovery than that of the groups that received a mixture of antibiotics. These changes could increase the risk of intestinal infections and promote inflammatory responses. Our findings suggest that patients who have been prescribed antibiotics should also receive supplementation with bacteria with beneficial potential to help restore the balance of their intestinal flora as soon as possible.

### Effects of long-term antibiotic use on the murine intestinal microbiota

In the experiments described above, we analyzed changes in intestinal microbiota diversity and structure induced by short-term antibiotic use. Long-term use of antibiotics exerts different selection pressure on bacteria, inducing drug resistance and leading to an increased in the abundance of opportunistic and pathogenic bacteria. Next, we analyzed the two long-term groups (male mice), which received either normal sterile water (control group) or continuous treatment with a mixture of antibiotics in their drinking water (100 mg/kg Ampicillin; 50 mg/kg Vancomycin; 100 mg/kg Metronidazole; and 100 mg/kg Neomycin) for 55 days (L-Mix group). Fecal samples were collected at 11 time points, and four of these were chosen for analysis: days 5, 15, 40, and 55. PCA analysis (Figure 4A) showed that the microbiota composition at all four time points from the control group clustered together, indicating that their composition was very similar. In contrast, the values for the four time points for the L-Mix group were scattered across the graph, indicating that there were relatively large differences in the microbiota composition between these four time points. This indicates that the impact of antibiotics on the intestinal microbiota is continuous, and that the microbiota structure is constantly changing. In addition, as shown in Figure 4B, at the phylum level, the abundance of *Bacteroidetes* (0.4754%, 0.2978%, 0.5595%) (D0, D40, D55) first decreased then increased. At the genus level (Figure 4C), the abundance of the bacteria with pathogenic potential *Bacteroides* (0.0468%, 0.1916%, 0.3461%) increased. We speculate that long-term antibiotic use led to the appearance of resistant strains in the intestinal microbiota. This effect will also apply to whole microbiomes, where the R/S ratios would be different from clade to another, as well as it’s natural the susceptibility to each antibiotic. As some susceptible species decrease in frequency or become extinct, new steady-state levels will be reached (David et al., 2019; Gjini and Brito, 2016).

**Figure 4.**
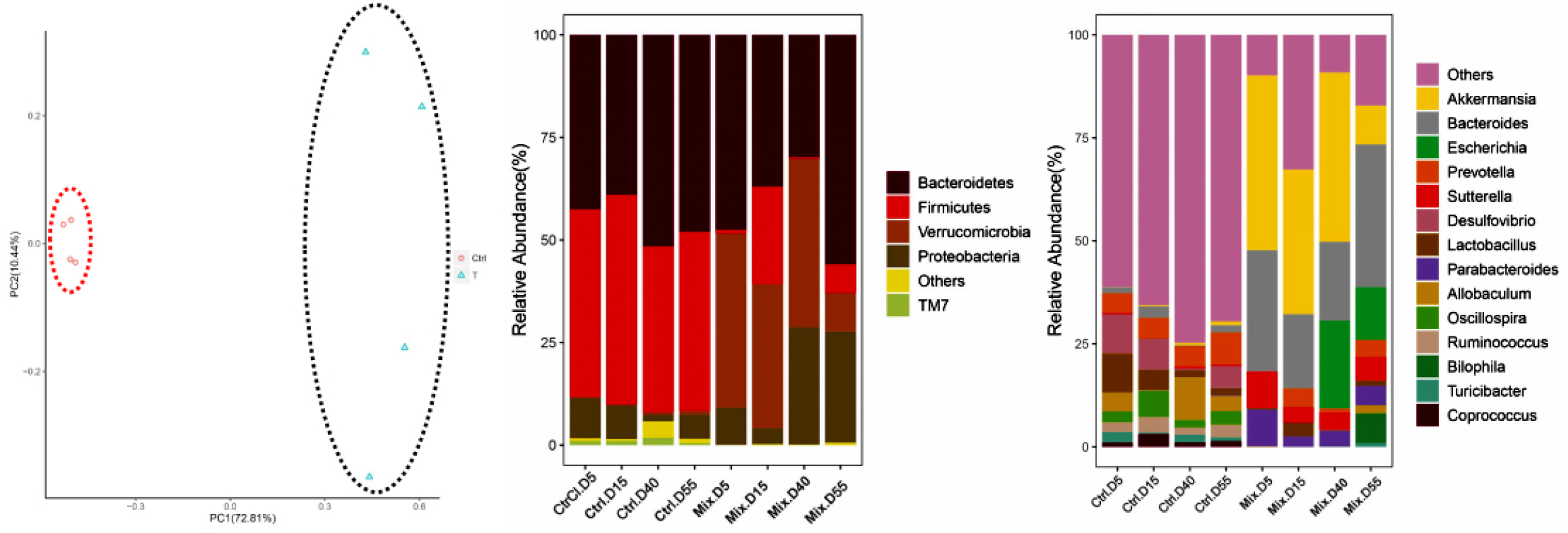
Effects of long-term antibiotic treatment on the composition of the murine intestinal microbiota. (A) PCA analysis. Ctrl: Control group, T: Mixture group. The more closely two samples group together on the graph, the more similar their composition. Values for samples taken from different treatment groups or environments can be either dispersed or aggregated, and the degree of dispersion can indicate the degree of similarity between the composition of samples taken under the same conditions. (B) and (C) show histograms of phylum and genus abundance, respectively. The sample names are shown on the abscissa, and the relative abundance of the annotated species is shown on the ordinate axis. Species whose abundance was less than 0.5% in all samples were merged into the ‘Others’ category.

## DISCUSSION

In this study, we used 16S rRNA gene sequencing technology to determine the short-term and long-term effects of Ampicillin, Vancomycin, Metronidazole, and Neomycin on the intestinal microbiota of mice by analyzing changes in the relative abundance of the bacteria with beneficial potential and the bacteria with pathogenic potential. We found that treatment with Ampicillin, Vancomycin, Metronidazole, Neomycin, and a combination of all four antibiotics decreased the diversity of the microbiota. Treatment with each of the four antibiotics resulted in a decrease in the abundance of bacteria with beneficial potential and an increase in the abundance of the bacteria with pathogenic potential. Previous studies have suggested that the decrease in the relative abundance of bacteria with beneficial potential observed after antibiotic use is due to a reduction in the absolute levels of bacteria with beneficial potential because of decreased microbiota diversity (Lankelma et al., 2017), and our results support this view. However, we believe that the changes that we observed in the intestinal microbiota reflect not only the antibiotics’ mechanisms of action, but also competition between different bacterial communities induced by intestinal microbiota dysbiosis. The decrease in the abundance of bacteria with beneficial potential is also affected by the increase in the abundance of the bacteria with pathogenic potential, and that this increase leads to a decrease in the relative abundance of the bacteria with beneficial potential. This increase in the abundance of the bacteria with pathogenic potential may also enhance the host inflammatory response and reduce the abundance of the bacteria with beneficial potential (Zárate-Bladés et al., 2016).

We also assessed the recovery of the murine intestinal microbiota 1 month after antibiotic treatment was stopped. In general, we found that the diversity of the intestinal microbiota recovered within a few weeks after discontinuation of antibiotic use, although the microbiota composition was different than before antibiotic treatment. Recent studies have found that, 4 months after stopping ciprofloxacin or clindamycin treatment, the abundance of only two OTUs had changed significantly, and the abundance of the other microbiota components had returned to baseline levels (Dethlefsen et al., 2008; Jakobsson et al., 2010). Our results are inconsistent with a previous study (Lankelma et al., 2017) because we found that, although the abundance of some bacterial genera recovered to baseline levels, the microbiota composition differed from that observed before antibiotic treatment, and in some cases these changes were permanent. Thus, our data indicate that antibiotics not only have a short-term effect on the host intestinal microbiota, but also a long-term effect. In addition to analyzing the effects of short-term use of the four antibiotics, we also analyzed the effects of long-term use of the four antibiotics on the intestinal microbiota. At the genus level (Figure 4C), the abundance of the opportunistic pathogen Bacteroides and of Escherichia increased. We speculate that long-term antibiotic used leads to the development of resistant strains, which is consistent with previous studies (Caballero et al., 2017; Hu et al., 2013; Kelly et al., 2020). In addition, our results show that pathogens such as Bacteroides and Escherichia are more likely to develop resistance.

Our study had some limitations. First, only three mice were included per group, but the changes in the intestinal microbiota caused by the antibiotics were drastic, so the results should be interpreted with caution. Second, the effect of antibiotics on the composition and function of the intestinal microbiota can change depending on the type and dosage of antibiotics used and the route of administration. In this study, we administered antibiotics orally to mice for 2 weeks, and did not explore the effects of different routes of administration on the murine intestinal microbiota. Third, we did not explore the mechanism underlying the changes we observed in the microbiota, which could be useful for designing strategies to restore normal host-microbiota interactions. Further research is needed to identify this mechanism.

In summary, our research shows that oral administration of four antibiotics, Ampicillin, Vancomycin, Metronidazole, and Neomycin, changes the composition of the murine intestinal microbiota, disrupts the balance between bacteria with beneficial potential and bacteria with pathogenic potential in the intestinal microbiota, and reduces the abundance of bacteria with beneficial potential and increases the abundance of the bacteria with pathogenic potential with short-term treatment. We speculate that these changes increase the risk of host infection. In addition, we found that oral antibiotics can have a long-term negative impact on the microbiota and produce drug-resistant bacteria, as indicated by the observation that, while the abundance of bacteria with beneficial potential recovered after antibiotics were withdrawn, the abundance of many bacteria with pathogenic potential did not return to baseline levels, and instead remained elevated. This effect is likely due to the host having been in a state of imbalance for a prolonged period of time. Use of these antibiotics should be considered carefully before being prescribed in the clinic, and restoration of the intestinal microbiota through fecal transplantation or probiotic administration should be considered.

## SEQUENCE DATA ACCESSION NUMBER

The original 16S rRNA sequence data are available at the NCBI by accession number PRJNA636223.

## AUTHOR CONTRIBUTIONS

L.H.L designed research; H.C, F.S.Y and H.F.J contributed to experimental work; H.C and F.S.Y performed the data analysis and wrote the manuscript; L.H.L. revised the manuscript.

## ACKNOWLEDGMENTS

The authors thank the support by the National Key Research and Development Program (2020YFC2002800), National Natural Science Foundation of China (Grant No 31671539), and Major Program of Development Fund for Shanghai Zhangjiang National Innovation Demonstration Zone<Stem Cell Strategic Biobank and Stem Cell Clinical Technology Transformation Platform> (ZJ2018-ZD-004).

## REFERENCES

Arias, C.A., and Murray, B.E. (2012). The rise of the Enterococcus: beyond vancomycin resistance. Nat Rev Microbiol 10, 266–278.

Bäumler, A.J., and Sperandio, V. (2016). Interactions between the microbiota and pathogenic bacteria in the gut. Nature 535, 85–93.

Buffie, C.G., Jarchum, I., Equinda, M., Lipuma, L., Gobourne, A., Viale, A., Ubeda, C., Xavier, J., and Pamer, E.G. (2012). Profound alterations of intestinal microbiota following a single dose of clindamycin results in sustained susceptibility to Clostridium difficile-induced colitis. Infect Immun 80, 62–73.

Caballero, S., Kim, S., Carter, R.A., Leiner, I.M., Sušac, B., Miller, L., Kim, G.J., Ling, L., and Pamer, E.G. (2017). Cooperating Commensals Restore Colonization Resistance to Vancomycin-Resistant Enterococcus faecium. Cell Host Microbe 21, 592-602.e594.

David, P.H.C., Sá-Pinto, X., and Nogueira, T. (2019). Using SimulATe to model the effects of antibiotic selective pressure on the dynamics of pathogenic bacterial populations. Biol Methods Protoc 4, bpz004.

Depommier, C., Everard, A., Druart, C., Plovier, H., Van Hul, M., Vieira-Silva, S., Falony, G., Raes, J., Maiter, D., Delzenne, N.M., et al. (2019). Supplementation with Akkermansia muciniphila in overweight and obese human volunteers: a proof-of-concept exploratory study. Nat Med 25, 1096–1103.

Dethlefsen, L., Huse, S., Sogin, M.L., and Relman, D.A. (2008). The pervasive effects of an antibiotic on the human gut microbiota, as revealed by deep 16S rRNA sequencing. PLoS Biol 6, e280.

Edgar, R.C. (2013). UPARSE: highly accurate OTU sequences from microbial amplicon reads. Nat Methods 10, 996–998.

Ferreira-Halder, C.V., Faria, A.V.S., and Andrade, S.S. (2017). Action and function of Faecalibacterium prausnitzii in health and disease. Best Pract Res Clin Gastroenterol 31, 643–648.

Freifeld, A.G., Bow, E.J., Sepkowitz, K.A., Boeckh, M.J., Ito, J.I., Mullen, C.A., Raad, II, Rolston, K.V., Young, J.A., and Wingard, J.R. (2011). Clinical practice guideline for the use of antimicrobial agents in neutropenic patients with cancer: 2010 update by the infectious diseases society of america. Clin Infect Dis 52, e56–93.

Getachew, B., Aubee, J.I., Schottenfeld, R.S., Csoka, A.B., Thompson, K.M., and Tizabi, Y. (2018). Ketamine interactions with gut-microbiota in rats: relevance to its antidepressant and anti-inflammatory properties. BMC Microbiol 18, 222.

Ghosh, S.S., Bie, J., Wang, J., and Ghosh, S. (2014). Oral supplementation with non-absorbable antibiotics or curcumin attenuates western diet-induced atherosclerosis and glucose intolerance in LDLR-/-mice--role of intestinal permeability and macrophage activation. PLoS One 9, e108577.

Gilbert, D.N., Moellering, R.C., and Eliopoulos, G.M. (2012). The Sanford guide to antimicrobial therapy 2012 (42nd Edition).

Gjini, E., and Brito, P.H. (2016). Integrating Antimicrobial Therapy with Host Immunity to Fight Drug-Resistant Infections: Classical vs. Adaptive Treatment. PLoS Comput Biol 12, e1004857.

González-Sarrías, A., Romo-Vaquero, M., García-Villalba, R., Cortés-Martín, A., Selma, M.V., and Espín, J.C. (2018). The Endotoxemia Marker Lipopolysaccharide-Binding Protein is Reduced in Overweight-Obese Subjects Consuming Pomegranate Extract by Modulating the Gut Microbiota: A Randomized Clinical Trial. Mol Nutr Food Res 62, e1800160.

Hagan, T., Cortese, M., Rouphael, N., Boudreau, C., Linde, C., Maddur, M.S., Das, J., Wang, H., Guthmiller, J., Zheng, N.Y., et al. (2019). Antibiotics-Driven Gut Microbiome Perturbation Alters Immunity to Vaccines in Humans. Cell 178, 1313-1328.e1313.

Han, Y., Yoon, J., and Choi, M.S. (2020). Tracing the Anti-Inflammatory Mechanism/Triggers of d-Allulose: A Profile Study of Microbiome Composition and mRNA Expression in Diet-Induced Obese Mice. Mol Nutr Food Res 64, e1900982.

Hu, Y., Yang, X., Qin, J., Lu, N., Cheng, G., Wu, N., Pan, Y., Li, J., Zhu, L., Wang, X., et al. (2013). Metagenome-wide analysis of antibiotic resistance genes in a large cohort of human gut microbiota. Nat Commun 4, 2151.

Jakobsson, H.E., Jernberg, C., Andersson, A.F., Sjölund-Karlsson, M., Jansson, J.K., and Engstrand, L. (2010). Short-term antibiotic treatment has differing long-term impacts on the human throat and gut microbiome. PLoS One 5, e9836.

Kelly, S.A., Rodgers, A.M., O’Brien, S.C., Donnelly, R.F., and Gilmore, B.F. (2020). Gut Check Time: Antibiotic Delivery Strategies to Reduce Antimicrobial Resistance. Trends Biotechnol 38, 447–462.

Kim, S.W., Suda, W., Kim, S., Oshima, K., Fukuda, S., Ohno, H., Morita, H., and Hattori, M. (2013). Robustness of gut microbiota of healthy adults in response to probiotic intervention revealed by high-throughput pyrosequencing. DNA Res 20, 241–253.

Lankelma, J.M., Cranendonk, D.R., Belzer, C., de Vos, A.F., de Vos, W.M., van der Poll, T., and Wiersinga, W.J. (2017). Antibiotic-induced gut microbiota disruption during human endotoxemia: a randomised controlled study. Gut 66, 1623–1630.

Le, J., Ngu, B., Bradley, J.S., Murray, W., Nguyen, A., Nguyen, L., Romanowski, G.L., Vo, T., and Capparelli, E.V. (2014). Vancomycin monitoring in children using bayesian estimation. Ther Drug Monit 36, 510–518.

Le Roy, T., Debédat, J., Marquet, F., Da-Cunha, C., Ichou, F., Guerre-Millo, M., Kapel, N., Aron-Wisnewsky, J., and Clément, K. (2018). Comparative Evaluation of Microbiota Engraftment Following Fecal Microbiota Transfer in Mice Models: Age, Kinetic and Microbial Status Matter. Front Microbiol 9, 3289.

Lewis, B.B., Buffie, C.G., Carter, R.A., Leiner, I., Toussaint, N.C., Miller, L.C., Gobourne, A., Ling, L., and Pamer, E.G. (2015). Loss of Microbiota-Mediated Colonization Resistance to Clostridium difficile Infection With Oral Vancomycin Compared With Metronidazole. J Infect Dis 212, 1656–1665.

Liévin-Le Moal, V., and Servin, A.L. (2014). Anti-infective activities of lactobacillus strains in the human intestinal microbiota: from probiotics to gastrointestinal anti-infectious biotherapeutic agents. Clin Microbiol Rev 27, 167–199.

Magoč, T., and Salzberg, S.L. (2011). FLASH: fast length adjustment of short reads to improve genome assemblies. Bioinformatics 27, 2957–2963.

Miao, Z., Lai, Y., Zhao, Y., Chen, L., Zhou, J., Li, C., and Lan, H. (2020). Scutellarein Aggravated Carbon Tetrachloride-Induced Chronic Liver Injury in Gut Microbiota-Dysbiosis Mice. Evid Based Complement Alternat Med 2020, 8811021.

Murgia, D., Angellotti, G., D’Agostino, F., and De Caro, V. (2019). Bioadhesive Matrix Tablets Loaded with Lipophilic Nanoparticles as Vehicles for Drugs for Periodontitis Treatment: Development and Characterization. Polymers (Basel) 11.

Muthuramalingam, K., Singh, V., Choi, C., Choi, S.I., Kim, Y.M., Unno, T., and Cho, M. (2020). Dietary intervention using (1,3)/(1,6)-β-glucan, a fungus-derived soluble prebiotic ameliorates high-fat diet-induced metabolic distress and alters beneficially the gut microbiota in mice model. Eur J Nutr 59, 2617–2629.

Perez-Lopez, A., Behnsen, J., Nuccio, S.P., and Raffatellu, M. (2016). Mucosal immunity to pathogenic intestinal bacteria. Nat Rev Immunol 16, 135–148.

Reid, G., Younes, J.A., Van der Mei, H.C., Gloor, G.B., Knight, R., and Busscher, H.J. (2011). Microbiota restoration: natural and supplemented recovery of human microbial communities. Nat Rev Microbiol 9, 27–38.

Rodrigues, R.R., Greer, R.L., Dong, X., KN, D.S., Gurung, M., Wu, J.Y., Morgun, A., and Shulzhenko, N. (2017). Antibiotic-Induced Alterations in Gut Microbiota Are Associated with Changes in Glucose Metabolism in Healthy Mice. Front Microbiol 8, 2306.

Sanders, M.E., Merenstein, D.J., Reid, G., Gibson, G.R., and Rastall, R.A. (2019). Probiotics and prebiotics in intestinal health and disease: from biology to the clinic. Nat Rev Gastroenterol Hepatol 16, 605–616.

Schloss, P.D., Westcott, S.L., Ryabin, T., Hall, J.R., Hartmann, M., Hollister, E.B., Lesniewski, R.A., Oakley, B.B., Parks, D.H., Robinson, C.J., et al. (2009). Introducing mothur: open-source, platform-independent, community-supported software for describing and comparing microbial communities. Appl Environ Microbiol 75, 7537–7541.

Seo, B., Jeon, K., Moon, S., Lee, K., Kim, W.K., Jeong, H., Cha, K.H., Lim, M.Y., Kang, W., Kweon, M.N., et al. (2020). Roseburia spp. Abundance Associates with Alcohol Consumption in Humans and Its Administration Ameliorates Alcoholic Fatty Liver in Mice. Cell Host Microbe 27, 25-40.e26.

Smits, W.K., Lyras, D., Lacy, D.B., Wilcox, M.H., and Kuijper, E.J. (2016). Clostridium difficile infection. Nat Rev Dis Primers 2, 16020.

Sommer, F., Anderson, J.M., Bharti, R., Raes, J., and Rosenstiel, P. (2017). The resilience of the intestinal microbiota influences health and disease. Nat Rev Microbiol 15, 630–638.

Sun, L., Zhang, X., Zhang, Y., Zheng, K., Xiang, Q., Chen, N., Chen, Z., Zhang, N., Zhu, J., and He, Q. (2019). Antibiotic-Induced Disruption of Gut Microbiota Alters Local Metabolomes and Immune Responses. Front Cell Infect Microbiol 9, 99.

Wang, K., Liao, M., Zhou, N., Bao, L., Ma, K., Zheng, Z., Wang, Y., Liu, C., Wang, W., Wang, J., et al. (2019). Parabacteroides distasonis Alleviates Obesity and Metabolic Dysfunctions via Production of Succinate and Secondary Bile Acids. Cell Rep 26, 222-235.e225.

Yang, J., Oh, Y.T., Wan, D., Watanabe, R.M., Hammock, B.D., and Youn, J.H. (2017). Postprandial effect to decrease soluble epoxide hydrolase activity: roles of insulin and gut microbiota. J Nutr Biochem 49, 8–14.

Zárate-Bladés, C.R., Horai, R., and Caspi, R.R. (2016). Regulation of Autoimmunity by the Microbiome. DNA Cell Biol 35, 455–458.

Zhang, C., Peng, Y., Mu, C., and Zhu, W. (2018). Ileum terminal antibiotic infusion affects jejunal and colonic specific microbial population and immune status in growing pigs. J Anim Sci Biotechnol 9, 51.

Zhang, Y., Liu, Q., Yu, Y., Wang, M., Wen, C., and He, Z. (2020). Early and Short-Term Interventions in the Gut Microbiota Affects Lupus Severity, Progression, and Treatment in MRL/lpr Mice. Front Microbiol 11, 628.

R Core Team (2018). R: A language and environment for statistical computing. R

